# Mixing of Porpoise Ecotypes in South Western UK Waters Revealed by Genetic Profiling

**DOI:** 10.1101/045807

**Authors:** Michaël C. Fontaine, Oliver Thatcher, Nicolas Ray, Sylvain Piry, Andrew Brownlow, Nicholas J. Davison, Paul Jepson, Rob Deaville, Simon J. Goodman

**Affiliations:** Groningen Institute for Evolutionary Life Sciences (GELIFES), University of Groningen, PO Box 11103 CC, Groningen, The Netherlands; School of Biology, Faculty of Biological Sciences. University of Leeds, LS2 9JT, UK; Institute of Zoology, Zoological Society of London, NW1 4RY, UK; Department of Zoology, University of Cambridge, CB2 3EJ, UK; EnviroSPACE Laboratory, Institute for Environmental Sciences, University of Geneva, Carouge, Switzerland; INRA, UMR CBGP, F-34988 Montferrier-sur-Lez Cedex, France; Scottish Marine Animal Stranding Scheme, SRUC Veterinary Services, Drummondhill, Stratherrick road, Inverness, IV2 4JZ UK.; Animal and Plant Health Agency, Polwhele, Truro, Cornwall, TR4 9AD, UK

**Keywords:** genetic admixture, continuous population, ecological genetics, dispersal, cetacean ecology, climate change, ecotype specialization

## Abstract

Contact zones between marine ecotypes are of interest for understanding how key pelagic predators may react to climate change. We analysed the fine scale genetic structure and morphological variation in harbour porpoises around the UK, at the proposed northern limit of a contact zone between southern and northern ecotypes in the Bay of Biscay. Using a sample of 591 stranded animals spanning a decade and microsatellite profiling at 9 loci, clustering and spatial analyses revealed that animals stranded around UK are composed of mixed genetic ancestries from two genetic pools. Porpoises from SW England displayed a distinct genetic ancestry, had larger body-sizes and inhabit an environment differentiated from other UK costal areas. Genetic ancestry blends from one group to the other along a SW-NE axis along the UK coastline, and showed a significant association with body size, consistent with morphological differences between the two ecotypes and their mixing around the SW coast. We also found significant isolation-by-distance among juveniles, suggesting that stranded juveniles display reduced intergenerational dispersal, while adults show larger variance. The fine scale structure of this admixture zone raises the question of how it will respond to future climate change and provides a reference point for further study.

## Introduction

Intraspecific differentiation in contiguous geographical areas due to vicariance or geographical barriers is common in nature^1^. However, in the marine environment, movements are typically unrestricted over vast distances for highly mobile species such as cetaceans. This raises the question of how populations become genetically and ecologically differentiated with eventual speciation^2^. Despite their high dispersal ability, some cetaceans show substantial population structure, sometimes over a small geographical scale, not necessarily associated with geographic distance^2–4^. In some cases, oceanographic processes and (or) behavioural traits explain a high level of population differentiation^4–9^. Prey availability, prey choice, social structure and/or other factors such as habitat availability, predator and competition pressure can all be involved in driving the pattern and extent of dispersal^3^. Explaining dispersal thus revolves around deciphering which current and/or historical mechanism(s) contributed to genetic structuring in the absence of obvious dispersal barriers.

The harbour porpoise (*Phocoena phocoena*) is one of the smallest and most abundant coastal cetaceans, widely distributed in sub-polar to temperate coastal waters of the northern hemisphere^10^. Numerous studies assessed the population genetic structure of harbour porpoises in the Western Palearctic waters (i.e. the eastern North Atlantic and Black Sea) during the last 20 years^4,11–15^. However, only recently have three ecotypes or subspecies been identified in Western Palearctic waters, based on genetic divergence of the mitochondrial genome, supported by morphological, and ecological differences^13^. These three ecotypes include an isolated population in the Black Sea (*P. p. relicta*), the southern ecotypes (*P. p. meridionnalis*) displaying larger body-size^16^, with two distinct populations inhabiting upwelling waters around Mauritania and the Iberian peninsula, and a northern ecotype (*P. p. phocoena*) inhabiting the continental shelf from the north side of the Bay of Biscay to the subarctic waters of Norway and Iceland.

Fontaine et al.^13^ showed that these 3 ecotypes resulted from an initial split between the North Atlantic and Mediterranean porpoises, with the colonization of the Mediterranean Sea during the last Ice Age. This event was followed by a split of the Mediterranean population into Eastern and Western groups from which descended the Black Sea population on one side^17^ and the Iberian and Mauritanian populations on the other side. Finally, the Iberian population came back into contact with the northern continental shelf ecotype during the last millennium, and most likely during the Little Ice age (*ca*. 600 years ago), establishing a contact zone on the northern side of the Bay of Biscay, with predominantly northward gene flow^13,18^. However, the fine scale spatial genetic structure of this admixture zone and the limits of its spatial distribution are still poorly understood. Previous studies had restricted sampling on the northern side of the Bay of Biscay, and in particular there has been limited coverage of porpoises from around the United Kingdom (UK).

In this study, we analysed the genetic structure of harbour porpoises around UK using a dense sampling of 591 stranded animals (Fig. 1 and electronic supplementary material (ESM), Fig. S1-S3) spanning a decade from 1990 to 2002 (ESM, Fig. S4). We test whether animals stranded around UK show any evidence of mixed genetic ancestry from distinct genetic pools and morphological differentiation in terms of relative body-size. Given the proximity of the Biscay admixture zone^4,13,18^, we should expect that porpoises in the SW part of UK would display evidence of such mixed ancestry and would display larger body size closer to Iberian porpoises. We also showed previously that gene flow and individual dispersal was restricted in space on the continental shelf North of the Bay of Biscay^4^, creating a pattern of Isolation by distance (IBD)^19,20^. Here, we test whether such IBD exists around UK and whether it differed between age classes. Understanding the physical and ecological factors which influence the distribution of different ecotypes is central to understanding how this key pelagic predator may react to future climate change, and its subsequent impacts on North East Atlantic ecosystem^21^.

**Figure 1.**
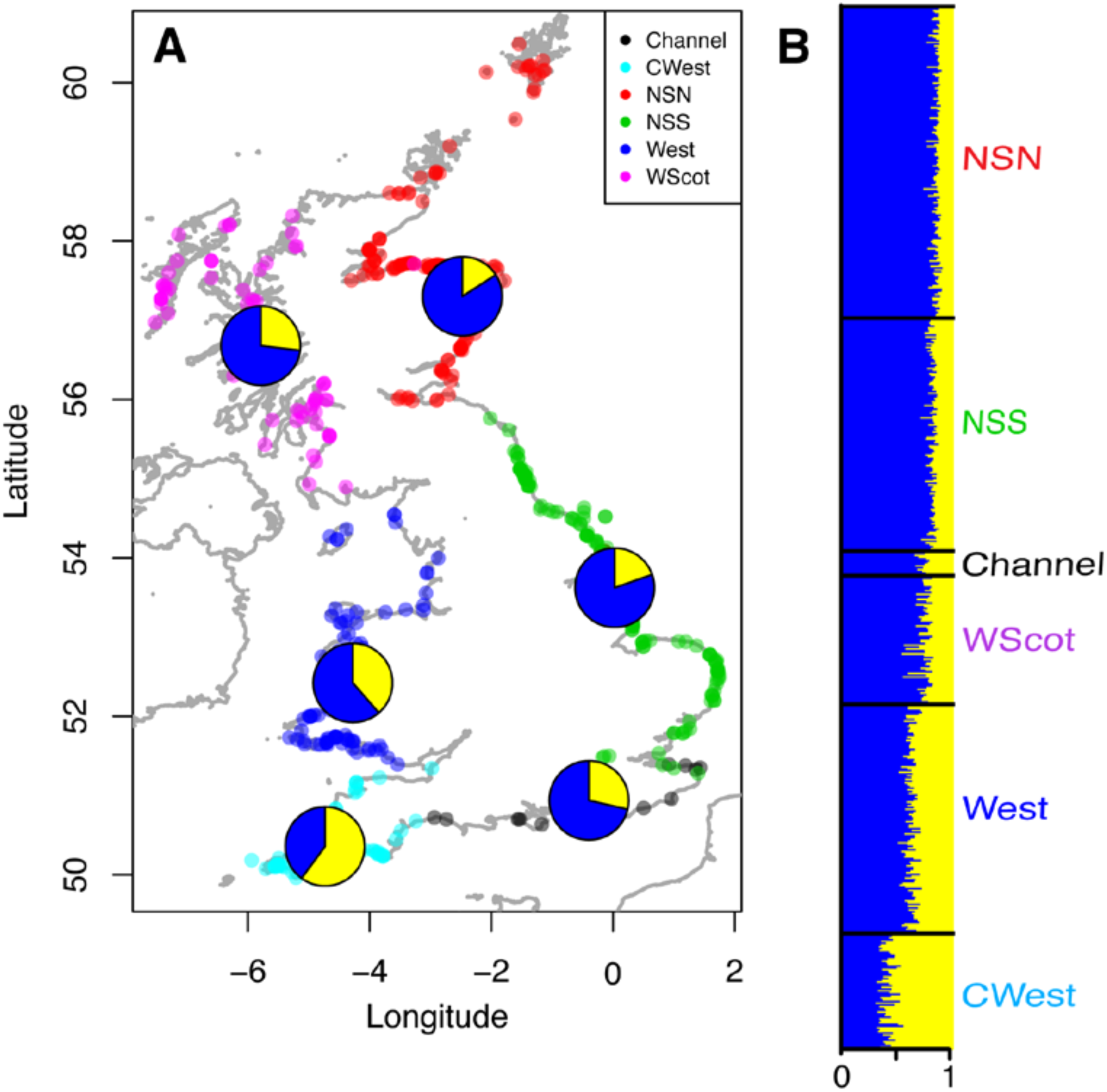
(*a*) Geographic locations of the harbour porpoises sampling (n=591) based on GPS coordinates or reported discovery location. Locations have been subdivided into 6 regions around UK and color-coded accordingly. Genetic structure of harbour porpoises in UK waters at K=2, as estimated by *Structure*, is displayed as the posterior admixture estimates averaged per regions. Panel (*b*) shows the individual admixture proportions. Each individual is represented by a column and the probability of that individual belonging to each cluster is indicated by coloured segments. Admixture proportions from *Structure* are based on the highest probability run (of ten) at that value of K=2.

## Material and Methods

### Sampling

Tissue samples collected between 1990 and 2002, body size, weight, age and associated temporal, geographical and life-history data for 591 stranded or by-caught porpoises from the United Kingdom Cetacean Strandings Project (http://ukstrandings.org/) archives were provided by P. Jepson (Institute of Zoology, Zoological Society of London) and R. Reid (Scottish Marine Animal Stranding Scheme, SRUC Veterinary Services, Inverness). The distribution of the sampling in space, time and per categories is shown in Fig. 1A, table 1, and ESM Fig. S1 to S4. All maps in this study were generated in R statistical environment v.3.0.226 using the shapefile from the UK coastline from European Commission, Eurostat (EuroGeographics – Countries 2014, CNTR_2014_03M_SH).

**Table 1.** Sampling distribution stratified by sex and age-class.

|  | **Females** | **Males** | **NA** | **Total** |
| --- | --- | --- | --- | --- |
| **Adult** | 86 | 109 | – | 195 |
| **Juvenile** | 126 | 117 | – | 243 |
| **Neonate** | 35 | 38 | – | 73 |
| **NA** | – | – | – | 80 |
| **Total** | 285 | 302 | 4 | 591 |

### Environmental data

Data on habitat characteristics across the study range with respect to salinity and sea surface temperature were taken from the National Oceanographic Data Centre (NODC) World Ocean Atlas (WOA01)^22^. Bathymetric data were extracted from the ETOPO2 dataset available on the US National Geophysical Data Centre (NGDC)^23^ and data on surface chlorophyll concentration were taken from the NASA Sea-viewing Wide Field-of-view Sensor database (SeaWIFS)^24^. To compare local habitat characteristics where harbour porpoise were living before dying, we calculated the mean value (± SD) of each variables within a radius arbitrarily set at 50km around each sampling locality using the Spatial Analyst extension in ArcGIS^™^ 8.2 (ESRI^®^).

### DNA extraction and microsatellite genotyping

Genomic DNA was extracted from skin or muscle sample using a standard phenol-chloroform protocol. Individuals were screened at 10 microsatellite loci used previously in harbour porpoises (Igf-1, 417/418, 415/416, GT011, GT136, GT015, EV94, EV104, GATA053, TAA031)^11^. PCR reactions were carried out in 10 μl volumes overlaid with 10 μl of mineral oil using 1 μl of template DNA (approximately 10-50 ng/*μ*l); 1 μl 10x PCR buffer with 1.5 mM MgCl_2_ (or 2.5 mM for loci GT015 and GT011, 2 mM for locus Igf-1), 45 nl Amplitaq DNA polymerase (Perkin Elmer), 0.8 mM of each primer, 0.1 mM of each nucleotide and 0.01 mM dCTP. PCR products were labeled by direct incorporation of <1*μ*Ci ^32^P-dCTP. The PCR cycle regime for EV104, EV96 and EV94 was: 1x (95°C for 3 minute); 7x (93°C for 1 minutes, 48°C for 1 minute, 72°C for 50 seconds); 25x (90°C for 45 seconds, second annealing temperature for 1 minute, 73°C for 1 minute); final stage (72°C for 15 minutes). For all other loci: 1x (3 minutes at 95°C); 35x (94°C for 1 minute, annealing temperature for 30 seconds, 72°C for 10 seconds); final stage (72°C for 15 minutes). PCR products from 96 individuals at a time were run on 6% denaturing polyacrylamide gels (Sequagel, National Diagnostics); visualisation was performed by autoradiography or Fujifilm BAS 2500 phosphor-imager. All the genotypes kept for analysis are consistent across 2 or more genotypings, and all homozygotes were rerun at lower annealing temperature to check for potential allelic drop out after initial analysis for Hardy Weinberg equilibrium on genotypes from the first screen.

### Data analysis

#### Genetic diversity and differentiation

We estimated the proportion of missing data per locus and region using *poppr* packages^25^ for the R statistical environment v.3.0.2^26^.

Observed and expected heterozygosity (*H*_*o*_, *H*_*e*_), allelic richness (*R*_*a*_), and inbreeding coefficient (*F*_*IS*_)^27^ were calculated using *GENETIX* v.4.05^28^ and *FSTAT* v.2.9.3^29^. These statistics were calculated per region (Fig. 1a). Per region *R*_*a*_ was computed based on a rarefaction procedure using the minimum sample size available across regions (n=13). We conducted permutation tests (10^5^ permutations) in *FSTAT* to assess potential departures from Hardy–Weinberg (HW) equilibrium for each population. Confidence interval at 95% for the *F*_*IS*_ values were calculated using the *diveRsity* v.1.9.89^30^ package for R^26^.

We also investigated local patterns of genetic diversity by calculating *R*_*a*_ on a grid lattice of 2° where cells included at least two samples. We used a custom R script to prepare the data, and *ADZE* 1.0^31^ to calculate *R*_*a*_ based on a standardized minimum sample size of 2 individuals. We plotted on a map an interpolated surface of *R*_*a*_ calculated using an inverse distance weighted procedure using *gstat* package for R^32^.

Levels of differentiation in allelic frequencies between regional groups of porpoises was estimated using pairwise *F*_*ST*_^27^ values and 95% confidence intervals (CIs) calculated using the *diveRsity*^30^ package for R. We considered *F*_*ST*_ comparisons as significant only if two conditions were met: the lower CI is >0, and P-values are <0.05 following a Bonferroni correction.

#### Bayesian genetic clustering analyses

We analysed the genetic structure using a Bayesian model-based clustering method implemented in *Structure* v.2.3.4^33–35^. Since in our dataset any genetic structure is likely to be due to weak IBD^4^, we introduced to the Bayesian analysis a *prior* assumption that individuals found in the same area are likely to be more closely related to each other than individuals sampled from more distant locations. To implement this, we used the sampling location as *a prior* information in the Bayesian inference using the *Locprior admixture* model^35^. This model has better performance to detect existing genetic structure when the level of divergence is weak, yet without introducing biases towards detecting structure when it is not present^35^. For that purpose, we divided the sampled area into 6 zones (Fig. 1A): the Channel, the Celtic Sea on the South West coast (CWest), the North Sea North (NSN), the North Sea South (NSS), the West coast (West), and West coast of Scotland (WScot), which correspond to the main maritime areas around the UK.

*Structure* analyses were conducted by running a series of independent simulations with different numbers of simulated clusters (*K*), testing all values from 1 to 5. Each run used an admixture model with correlated allele frequencies, 1 × 10^6^ iterations after a burn-in of 1 × 10^5^ iterations. Ten replicates of each run were conducted to test for convergence of the MCMCs. *Structure* results were then post-processed using *StructureHarvester*^36^ and plotted using custom R-scripts.

Since our present data set does not include samples from the southern ecotypes along the Iberian waters for a full set of loci compatible with previous studies, we tested empirically the impact of omitting this group on the clustering solution identified by *Structure*. For that purpose, we used data previously published by Fontaine *et al*.^13^ focusing on the area surrounding the Bay of Biscay, including individuals from the southern ecotype along the Iberian coasts, from the admixed zone in the northern Bay of Biscay, Irish seas, Scotland, Channel, and from the northern ecotypes along the Belgian and Dutch coasts. We run *Structure* using the same conditions as described above and, including or not the Iberian individuals.

#### Non-parametric multivariate analyses

Multivariate analyses of genetic data can provide a complementary view to model-based Bayesian clustering approach^37,38^. We further analysed the genetic structure using a Principal Component analyses (PCA)^38–40^ and a modified version of this analysis, known as spatial PCA or sPCA^41^, accounting for spatial autocorrelation, and aiming at displaying genetic variance with a spatial structure. We used a ‘global’ and ‘local’ test procedures based on Monte Carlo permutations (10^4^ permutations) to interpret the significance of the spatial principal components in the sPCA^41^. Following the definition of the sPCA, ‘global structure’ relates to patterns of spatial genetic structure, such as patches, clines, isolation by distance and intermediates, whereas ‘local structure’ refers to strong differences between local neighbourhoods^41^. These analyses were conducted using *adegenet 1.4-2* package^42^ for the R software^26^.

#### Isolation by distance analysis

Patterns of isolation by distance (IBD) may emerge if dispersal is spatially restricted at the scale of our study^19^. Under the hypothesis of IBD, genetic differentiation between individuals (estimated using the *â*_*r*_ statistics analogous to *F*_*ST*_/(1-*F*_*ST*_) between demes) is expected to increase with increasing geographic distance^20,43,44^. We calculated the regression coefficient (b) between genetic and geographic distance matrices between individuals and evaluated its significance with a Mantel Test (10^4^ permutations of geographic locations) using *SpaGeDi* 1.4^45^. Instead of using an Euclidian distance that would poorly describe the actual geographic distance between pairs of individuals, we computed a marine geographic distances that accounts for the shortest path by sea between two individuals as described in Fontaine *et al*.^4^. To compute this marine geographic distance, we used a Least Cost Path algorithm using a modified version of PATHMATRIX^46^, implemented in *C* for improved computational efficiency (available upon request to N Ray).

We tested the occurrence of IBD first on the entire data set. Then we tested whether IBD patterns differed among sex and age classes. While IBD patterns should be similar among sexes, since we are using autosomal loci, IBD patterns can potentially differ among age classes if one of the classes (e.g., the juveniles) disperses less than another classes (e.g. the adults). We tested IBD in adults versus juveniles only, as sample sizes (table 1), spatial (ESM Fig. S1 and S2) and temporal distributions (ESM Fig. S4), were not sufficient to partition the data further and maintain satisfactory statistical power.

#### Morphological analysis

Data on body-length, age and sex was available for a large subset of the individuals (n=336) included in the genetic analyses. As the two porpoise ecotypes are likely present in the studies area and are known to differ according to their body size^16^, we investigated how body-length varied as a function of the animal age and sex using a linear model. We were particularly interested in the residual variation not accounted for by the age and sex and in particular its geographic component. Residual variation in body length was compared among the 6 geographic zones with an ANOVA in R^26^ using log-transformation for the body-length and the age. We also assessed the correlation between individual residual body size and individual admixture score derived from the *Structure* analysis.

## Results

### Genetic diversity and differentiation between regions

The proportion of missing data observed at the 10 microsatellite loci ranged between 0.5% and 4.9% (ESM Fig. S5). All loci but EV104 showed less than 10% missing data in any of the 6 geographic regions around UK (Fig. 1a and ESM Fig. S5). We excluded locus EV104 from further analyses as the proportion of missing data exceeded 10% in some regions (ESM Fig. S5) and potential null alleles have been recorded in other studies^4^. The genetic diversity (also known as expected heterozygosity, *H*_*e*_) of the remaining 9 loci is shown in table 2, and ranges between 0.67 and 0.71 with an average value of 0.69 ± 0.01 across the six regions. The allelic richness per region ranged between 3.3 and 3.7 alleles for a standardized sample size of 13 individuals. Overall, none of the loci displayed any significant departure from Hardy-Weinberg and Linkage Equilibrium expectations.

**Table 2.**
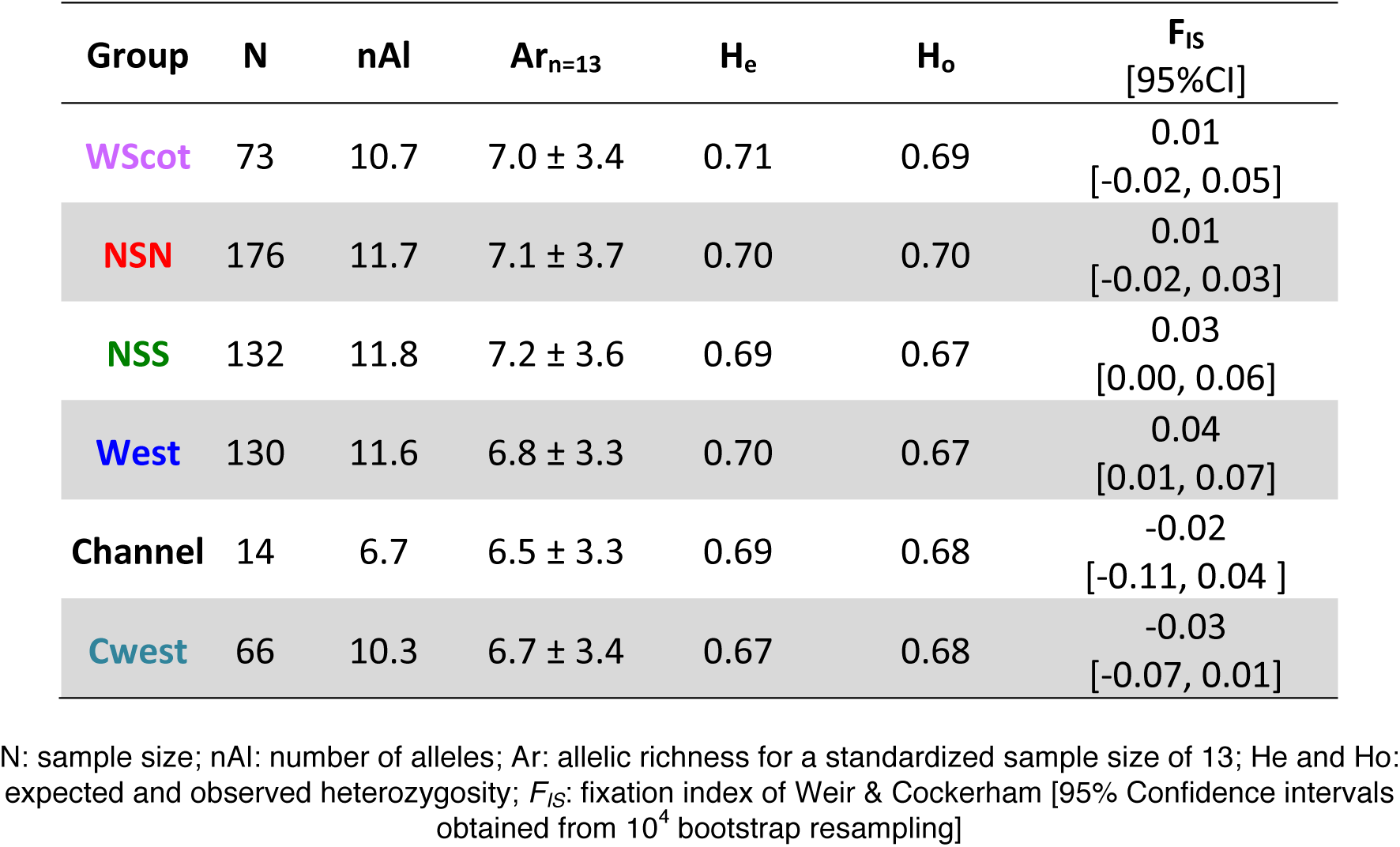
Genetic variation at the 9 microsatellite loci.

### Genetic structure

Differences in allelic frequencies estimated using *F*_*ST*_ between porpoises from the six regions ranged between 0.0 and 1.3% (table 3). Only the Cwest group display consistently small but significant *F*_*ST*_ values when compared to porpoises from the five other geographic regions.

**Table 3.** *F*_*ST*_ value [95% confidence intervals estimated using 10^4^ bootstrap resampling] (below) and P-value estimated using 10^4^ permutations (above). In bold are the pairwise comparisons that are statistically significant after a Bonferroni correction at *a*=0.05 and with a low 95% CI > 0.

| FST | Channel | Cwest | NSN | NSS | West | Wscot |
| --- | --- | --- | --- | --- | --- | --- |
| Channel | - | 0.010 | 0.090 | 0.047 | 0.113 | 0.333 |
| Cwest | 0.013<br>[-0.007, 0.043] | - | **0.003** | **0.003** | **0.003** | **0.003** |
| NSN | 0.006<br>[-0.010, 0.028] | **0.012**<br>**[0.006, 0.020]** | - | 0.523 | **0.003** | 0.017 |
| NSS | 0.006<br>[-0.009, 0.029] | **0.010**<br>**[0.004, 0.017]** | 0.001<br>[-0.002, 0.004] | - | 0.010 | 0.033 |
| West | 0.002<br>[-0.012, 0.023] | **0.008**<br>**[0.001, 0.016]** | **0.003**<br>**[0.000, 0.007]** | 0.003<br>[0.000, 0.007] | - | 0.007 |
| Wscot | 0.001<br>[-0.014, 0.023] | **0.012**<br>**[0.004, 0.022]** | 0.000<br>[-0.003, 0.004] | 0.003<br>[-0.002, 0.009] | 0.002<br>[-0.002, 0.007] | - |

The Bayesian clustering analyses in *Structure* (Fig. 1) also showed that porpoises from the South West (CWest) were genetically differentiated from the other groups, with an admixed genetic ancestry (in yellow), the proportion of which in the genetic pool, progressively declines from along a SW-NE axis, deeper into the Channel and Irish Sea (Fig. 1b).

The spatial Principal Component Analysis (sPCA) provides a similar picture of the genetic structure (Fig. 2). The Global test assessing the significance of positive sPCs showed that the first sPC is indeed significant (*p* = 0.004) and support the existence of a global genetic structure such as cline or clusters^41^. In contrast, the local test showed that none of the negative sPC are significant (*p* = 0.598). Plotting the individual scores along the first two positive sPCs (Fig. 2a) shows that porpoises from the SW region of UK (Cwest) depart from the others along the first sPC axis and that the genetic composition of British porpoises gradually change along a SW-NE geographic axis (Fig. 2a). This spatial structure is also very well depicted when plotting individual scores for the sPC1 on a map (Fig. 2b).

**Figure 2.**
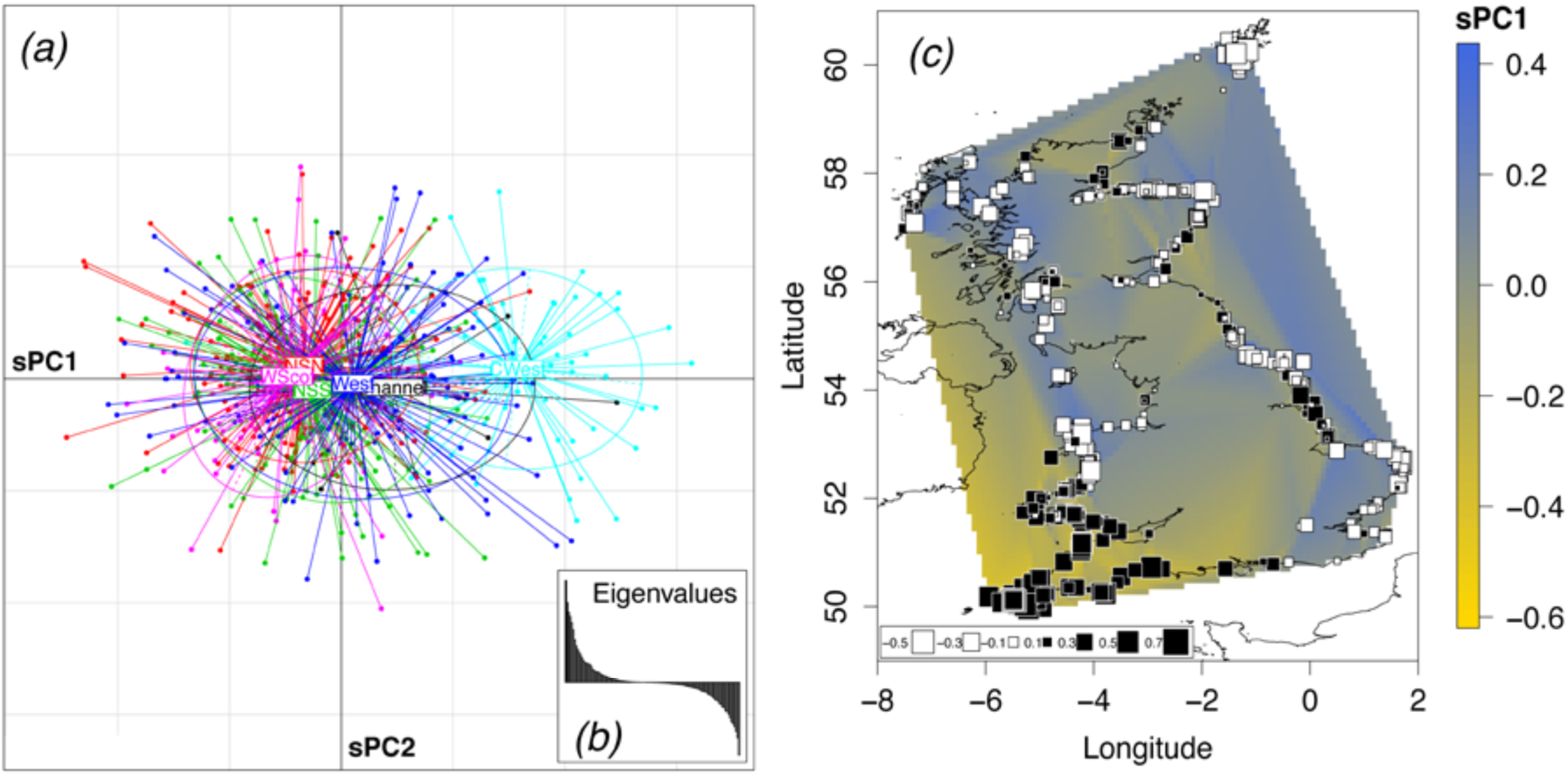
Spatial principal component analysis (sPCA). (*a*) The scores for each individual genotype are plotted for the first two sPCs, with colors indicating the discovery localities (see Fig. 1). (*b*) The inset provides the positive and negative eigenvalues. (*c*) Individual scores for the first component of the sPCA are displayed on the map using a size gradient of squares and a spatial interpolation surface.

The most parsimonious explanation for the distinct genetic composition of the SW porpoises is that it arises from the admixture with the southern ecotype from Iberian waters^13^. Excluding samples from Iberian porpoises did not affect the ability of the *Structure* algorithm to recover the correct genetic structure in a reanalysis of a previously published data set^13^ (ESM, Fig. S6), yielding signatures of structure and admixture consistent with those observed in the current study.

#### Isolation by distance (IBD)

We found significant IBD throughout the whole sample, indicating that gene flow, and thus individual dispersal, is spatially restricted (table 4). The IBD slope was similar between males and females, as expected since we are analysing autosomal loci. When structuring by age-class, the IBD slope was 4 times higher in juveniles than in adults, only being significantly greater than zero in the former. This suggests that juveniles mostly drive the IBD signal, while adults display a higher variance in dispersal pattern.

**Table 4.**
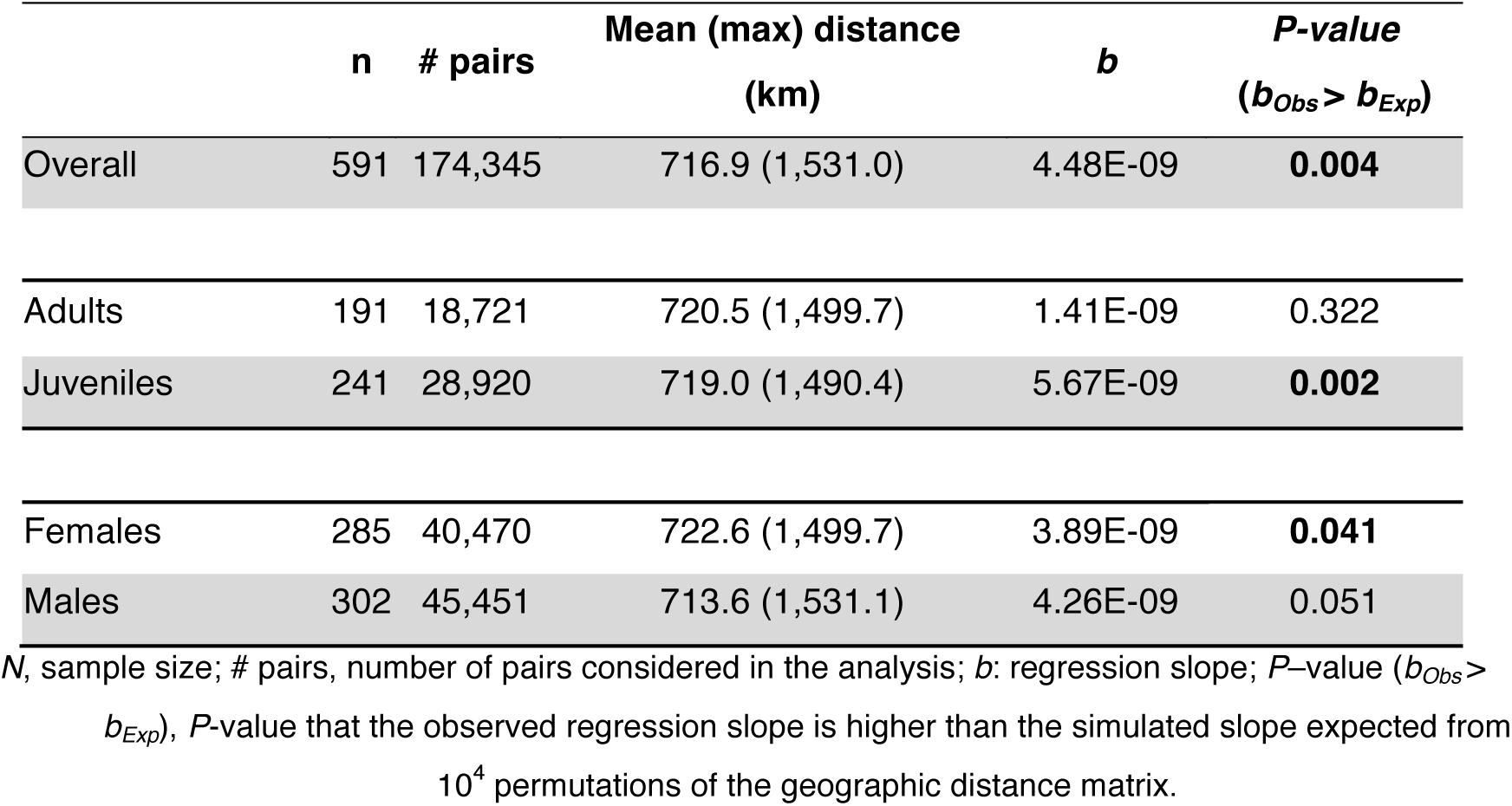
Isolation by distance conducted at individual level between porpoises.

#### Morphological analyses

As previously reported^47^, we found that both age and sex were significant predictors of the body-length, explaining about 61% of the total variation (Linear model, LM1: *F*_2,334_ = 261.1, *p* < 2.2 10^-16^, n=336). We inspected the geographic variation in the residuals (Fig. 3a and 3b) and observed that porpoises from the SW (Cwest) area as well as some porpoises from the West area of England displayed significantly larger body-size compare to the others (one-way ANOVA, *F*_5_ = 15.53, *p* < 9.9 10^-14^ and *p* < 0.001 for all Tukey pairwise comparisons involving Cwest, table S1). We also observed a strong correlation between individual residuals of body size and individual genetic admixture proportions estimated in the Bayesian clustering analysis of *Structure* (Pearson *r* = 0.39, *p* = 8.3 10^-14^, Fig. 3c). Combining the genetic ancestry together with the age and sex in the linear model for predicting the body length increased significantly the total variance explained by the linear model up to 67% (LM2: *F*_3,333_ = 225.5, *p* < 2.2 10^-16^). This model with genetic ancestry offered a significantly better fit to the data compare to a model where it is not included (nested model comparison LM1 *vs* LM2: ANOVA *F*_1,333_ = 60.8, *p* < 8.2 10^-14^).

**Figure 3.**
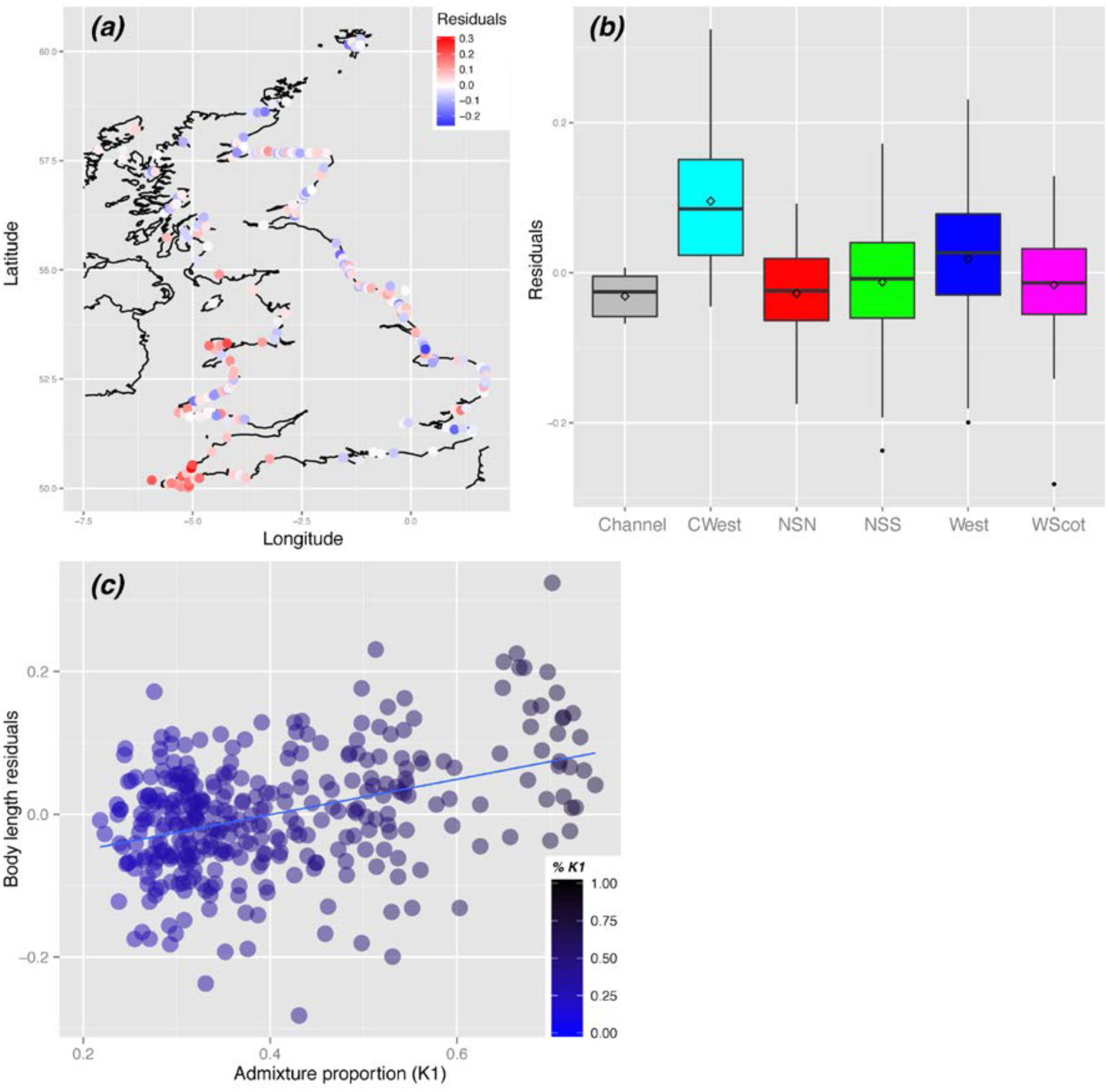
Geographical variation in the residuals from the linear model of the body-length values as a function of the age and sex. (*a*) Residual values are shown on a map and (*b*) as boxplots per regions. Panel (*c*) shows the relationship between the individual residuals of body size with individual genetic admixture proportions (%*K1*) estimated in the Bayesian clustering analysis of *Structure* (Pearson *r* = 0.39, *p* = 8.3 10^-14^).

## Discussion

Harbour porpoises in UK waters are part of a genetic continuum, characterized by a weak genetic structure, in which geographically proximate individuals are genetically more similar, a so called isolation by distance (IBD) pattern^4,13^. However, porpoises stranded along the SW coasts of the UK, facing the Celtic Sea and the Atlantic side of the Channel, display significant genetic differentiation (Fig. 1 and 2), which appear to be driven by admixture with more southerly populations, since the extent of differentiation gradually changes along a SW-NE axis moving deeper into the English Channel and Irish Sea. The genetic distinctiveness of these SW porpoises, shown independently by pairwise *Fst* comparisons (table 2), the Bayesian clustering analysis (Fig. 1) and the sPCA (Fig. 2), is coincident with their significantly larger body sizes compared to the rest of the UK, reminiscent of the larger body size of the southern ecotypes inhabiting the coastal Atlantic waters of Iberia^13,16^. In addition, we observed a significant correlation between body size and admixture proportion, suggesting a strong link between genotype and morphology throughout the porpoise distribution around the UK. This represents the largest assessment of body size variation in European porpoises to date, and to our knowledge the first report of a potential association between genotype and body size variation at a population level in a cetacean.

It was not possible to incorporate genetic data from previous studies^13^, with individuals from across the whole eastern Atlantic range directly into the current work, due to insufficient overlap of microsatellite loci employed. Therefore, the source populations that might contribute to the admixture signal seen in the SW UK cannot be directly identified here. However, admixture between southern and northern ecotype was previously detected in the northern side of the Bay of Biscay, the Celtic Sea and in the western side of the Channel on a sampling of the same period of time in Fontaine *et al*.^13^. Therefore, the most parsimonious explanation for both the genetic and morphological variation around UK coastline is that stranded porpoises along the SW coasts are primarily composed of individuals having an admixed ancestry, forming the northern tip of the contact zone between the northern ecotypes inhabiting the continental shelf and the southern ecotype which inhabits the coastal Atlantic waters of the Iberian peninsula^13^, with a gradual transition between the two. The previous studies had only relatively sparse sampling along the French Channel and Irish coasts, with none from the SW UK, so the current analysis defines the limits of the admixture zone more precisely and shows how it extends through the Channel and Irish Sea. The local marine environment where porpoises from CWest area were living before dying also differed substantially from the other regions with waters that are warmer, saltier, slightly lower surface Chlorophyll concentration on average (Fig. 4). This area encompassing the Celtic Sea corresponds to the transition between two biogeographic marine zones (the Boereal-Lusitanean transition following^48^), the warm-temperate waters and cool-temperate waters^49^.

**Figure 4.**
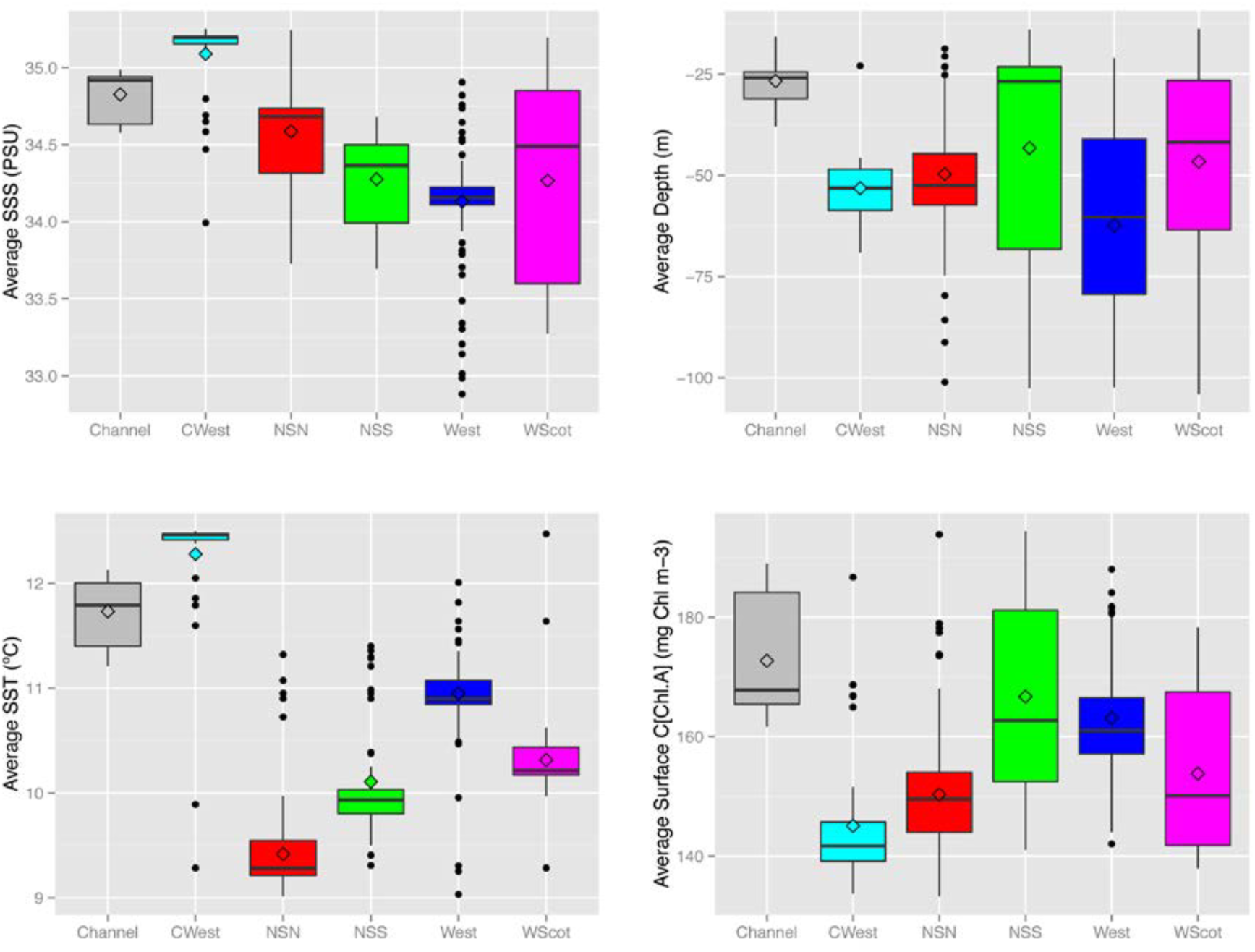
Box plot describing the environment along the UK coastline within a 50km radius surrounding stranded harbour porpoises. Annual Sea Surface Salinity (SSS), Temperature (SST), Depth, and Sea Surface Chlorophyll concentration are shown.

Interestingly, porpoises from SW coasts facing the northern part of the Bay of Biscay displayed slightly lower genetic diversity compared to more northern porpoises (ESM Fig. S7). A previous genetic study reported a similar pattern at larger scale in the Bay of Biscay together with a stronger IBD pattern than in the North Sea (see table 2 in Fontaine et al.^4^ and table S8 in Fontaine et al.^13^). Such reduced genetic diversity in a zone of admixture might appear counter intuitive at first glance, since we would usually expect an increase of genetic diversity when two distinct populations meet in a contact zone. However, the previous studies showed that genetic diversity of the Iberian population is very low and does not have any private alleles relative to the northern continental shelf populations. Therefore the reduction in diversity of the Biscay contact zone could arise through a combination of low genetic diversity of the southern ecotypes, and a high level of unidirectional gene flow from the Iberian population to the northern populations^4,13,18^. This results in a smaller effective population size and stronger IBD slope, which is inversely related to the product of local effective population size and the neighbourhood size (i.e. squared variance of the intergenerational dispersal distance)^20,43,44^.

The isolation by distance observed around the UK was weak but highly significant, and consistent with patterns observed in other parts of the range^4^. We did not observe any differences in IBD slope between sexes. However, we observed that juveniles displayed an IBD slope four times steeper than adults. While distinct local effective population size or genetic diversity cannot explain this difference, the likely explanation is that juveniles may show a reduced intergenerational dispersal distance compared to adults. Adults have the time and opportunity to disperse further away from their birthplaces than juveniles. The intergenerational dispersal distance and especially its variance component should thus be reduced in juveniles relative to adults, as suggested by our results.

The evidence of an admixed contact zone between northern and southern porpoise ecotypes, extending from the northern Bay of Biscay to waters around the SW United Kingdom, identified in this and previous studies^13^, raises the question of what environmental and ecological factors determine the distributions of the ecotypes, extent of the contact zone, and whether the distributions are stable or dynamic. Previous work has shown that the structure and distribution of harbour porpoise populations has been influenced by changes in oceanographic conditions which affect food resources ^4, 13^. Therefore the location and extent of the Biscay admixture zone is likely to be similarly dynamic and sensitive to past and future changes in climate which influence shifts in oceanographic and ecological conditions. For instance warming waters may see a northward expansion of the southern ecotype, which would be detectable by a shift in the extent of the admixture zone around the SW United Kingdom. The data presented here represent samples spanning an approximate 12 years window from 1990-2002. Future genetic studies, making use of the now extensive time series of samples spanning several decades available from European cetacean stranding programmes, will help test whether contemporary porpoise populations are showing a dynamic response to current climate change, and could be important in understanding how the structure of European marine ecosystems might respond to changes in the populations of such keystone predators^21^.

## Acknowledgement

We thank Prof. William Amos, Department of Zoology University of Cambridge, for providing laboratory facilities and other support for the microsatellite genotyping work. We also thank Bob Reid for the provision of the samples from Scotland.

## Author contribution

MCF and SJG designed the study; PJ, RD and ND collected samples and conducted necropsies; OT performed the laboratory experiment and data collection; MCF analysed the data with help from NR and SP; MCF wrote the manuscript with help from SJG and approval by all co-authors.

## Additional information

### Data accessibility

All data underlying this publication has been made publicly available in Dryad [to be provided upon acceptation of the paper].

### Funding

Collection and curation of porpoise tissue samples examined in this research were collected under the aegis of the collaborative Cetacean Strandings Investigation Programme, which is funded by the Department for Environment, Food and Rural Affairs (Defra) and the Devolved Governments of Scotland and Wales, as part of the UK government’s commitment to a number of international conservation agreements. OT and laboratory costs were supported by a Natural Environment Research Council (NERC) PhD studentship held by the Institute of Zoology and Department of Zoology University of Cambridge (ref: NER/S/A/2001/06405). MCF was supported by a short-term Marie-Curie Fellowship from the AGAPE program (Univ. of Leeds, UK; ref: MEST-CT-2004-504318).

### Competing Financial Interests

The authors declare no competing financial interests.

